# Transcriptional analysis of peripheral memory T cells reveals Parkinson’s disease-specific gene signatures

**DOI:** 10.1101/2021.05.28.446215

**Authors:** Rekha Dhanwani, João Rodrigues Lima-Junior, Ashu Sethi, John Pham, Gregory Williams, April Frazier, Yaqian Xu, Amy W. Amara, David G. Standaert, Jennifer G. Goldman, Irene Litvan, Roy N. Alcalay, Bjoern Peters, David Sulzer, Cecilia S. Lindestam Arlehamn, Alessandro Sette

## Abstract

Parkinson’s disease (PD) is a multi-stage neurodegenerative disorder with largely unknown etiology. Recent findings have identified PD-associated autoimmune features including roles for T cells. To further characterize the role of T cells in PD, we performed RNA sequencing on PBMC and peripheral CD4 and CD8 memory T cell subsets derived from PD patients and age-matched healthy controls. When the groups were stratified by their T cell responsiveness to alpha-synuclein (α-syn) as a proxy for ongoing inflammatory autoimmune response, the study revealed a broad differential gene expression profile in memory T cell subsets and a specific PD associated gene signature. We identified a significant enrichment of transcriptomic signatures previously associated with PD, including for oxidative stress, phosphorylation, autophagy of mitochondria, cholesterol metabolism and inflammation, and the chemokine signaling proteins CX3CR1, CCR5 and CCR1. In addition, we identified genes in these peripheral cells that have previously been shown to be involved in PD pathogenesis and expressed in neurons, such as LRRK2, LAMP3, and aquaporin. Together, these findings suggest that features of circulating T cells with α-syn-specific responses in PD patients provide insights into the interactive processes that occur during PD pathogenesis and suggest potential intervention targets.

## Introduction

Parkinson’s disease (PD) is a progressive neurodegenerative disorder characterized by two hallmarks: (i) loss of dopaminergic neurons in the substantia nigra (SN) of the brain responsible for the motor features (Fahn and Sulzer, 2004) and (ii) excess accumulation of aggregated α-synuclein (α-syn) protein (Spillantini et al., 1997). This loss of dopaminergic neurons in the SN is believed to be the reason for the parkinsonian motor signs (increased rigidity, slowness, rest tremor, and at later stages postural instability) observed in PD (Archibald et al., 2013). There are approximately 1 million people in North America affected with this debilitating disease (Marras et al., 2018). The diagnosis and management of PD is challenging as the disease is constrained by limited treatment options, which are mainly focused on improving postural instability and non-motor (constipation, mood, sleep, cognition) symptoms. Considering the increasing prevalence and overall societal impact of PD, it is imperative to explore the underlying mechanisms that play a role in the progression of this heterogenous and complex disease and ultimately to develop targeted symptomatic and disease-modifying interventions.

Several lines of evidence highlight an association of PD with inflammation. In 1988, a landmark postmortem study by McGeer and colleagues reported activated microglia in SN of PD subjects (McGeer et al., 1988). Since then, several reports and studies have indicated an association between an enhanced inflammatory response and PD (Stojkovska et al., 2015).

Recent studies have revealed an autoimmune component in PD, which comprises recognition of several α-syn-derived T cell epitopes by CD4 T cells (Sulzer et al., 2017), and demonstrate an increased α-syn-specific T cell reactivity in preclinical and early stages of the disease (Lindestam Arlehamn et al., 2020).

These observations can be interpreted in the broader context of current understanding of PD pathogenesis and progression. It is widely thought that clinically diagnosed motor PD and cognitive impairment is preceded by a long (often decades) prodromal phase, associated with symptoms ranging from alteration of sense of smell, constipation and sleep disorders that may precede the loss of SN dopaminergic neurons (Postuma and Berg, 2016). Indeed, in a single case study where T cell samples were available years prior to and after disease onset, α-syn-specific CD4 T cells were detected at higher levels before disease symptomatic onset (Lindestam Arlehamn et al., 2020).

To define molecular alterations associated with PD, we compared the transcriptional profiles of peripheral T cells derived from individuals with diagnosed motor PD to those of age-matched healthy controls (HC). We hypothesized that PD patients exhibiting α-syn-specific T cell responses (PD Responders; PD_R) are associated with an inflammatory stage of the disease, while the non-responder category (PD_NR) is associated with a non-α-syn-specific and/or later stage when inflammatory features of the disease have subsided. We have identified differences in transcriptomic profiles associated with both CD4 and CD8 T cell subsets that are apparent if the PD subjects are classified based on T cell reactivity to α-syn (PD_R vs. PD_NR). The results indicate that genes differentially regulated in CD4 memory T cells were enriched in oxidative stress and autophagy functions, while those upregulated in CD8 memory T cells were enriched in inflammatory and chemotaxis-related gene functions.

## Results

### Classification of PD subjects based on α-syn specific T cell reactivity

In previous studies, we detected α-syn specific T cell responses in approximately 40-50% of PD subjects (Lindestam Arlehamn et al., 2020; Sulzer et al., 2017). We further reported that α-syn specific T cell reactivity is specifically associated with preclinical and early timepoints following onset of motor PD features (Lindestam Arlehamn et al., 2020), while responses subsided in later stages of PD. Based on this finding, we hypothesized that PD subjects that demonstrate α-syn-specific T cell reactivity could be a “proxy” for individuals associated with an active inflammatory autoimmunity phenotype, and that analysis might reveal a transcriptional profile distinct from subjects without PD (healthy controls; HC) or PD subjects that do not exhibit α-syn T cell reactivity.

Accordingly, based on the magnitude of total response mounted against α-syn peptides, PD subjects were classified in two categories: responders (denoted as PD_R; > 250 SFC for the sum of IFNγ, IL-5, and IL-10) and non-responders (denoted as PD_NR; < 250 SFC). We also included age-matched HC who were α-syn non-responders (HC_NR), to avoid the possibility that HC who exhibit α-syn-specific T cell reactivity may be in prodromal stages of PD. The classification criteria were based on our previously published studies (Lindestam Arlehamn et al., 2020; Sulzer et al., 2017) where we determined α-syn-specific T cell reactivity for PD following *in vitro* restimulation assays, and measured cytokine release by Fluorospot or ELISPOT assays.

To investigate differential gene expression signatures, we examined 34 PD subjects including PD_R (n=14) and PD_NR (n=20) (**Figure 1A**). For control subjects, we selected 19 HC_NR subjects. We first analyzed the relative frequency of major PBMC subsets, i.e., monocytes, NK cells, B cells, T cells, and CD4 and CD8 memory T cells by flow cytometric analysis. The frequency of each PBMC subset was remarkably similar in all groups (**Supplementary Figure 1A**) and there was no significant difference between CD4 and CD8 memory T cell subsets (**Supplementary Figure 1B-C**).

**Figure 1.**
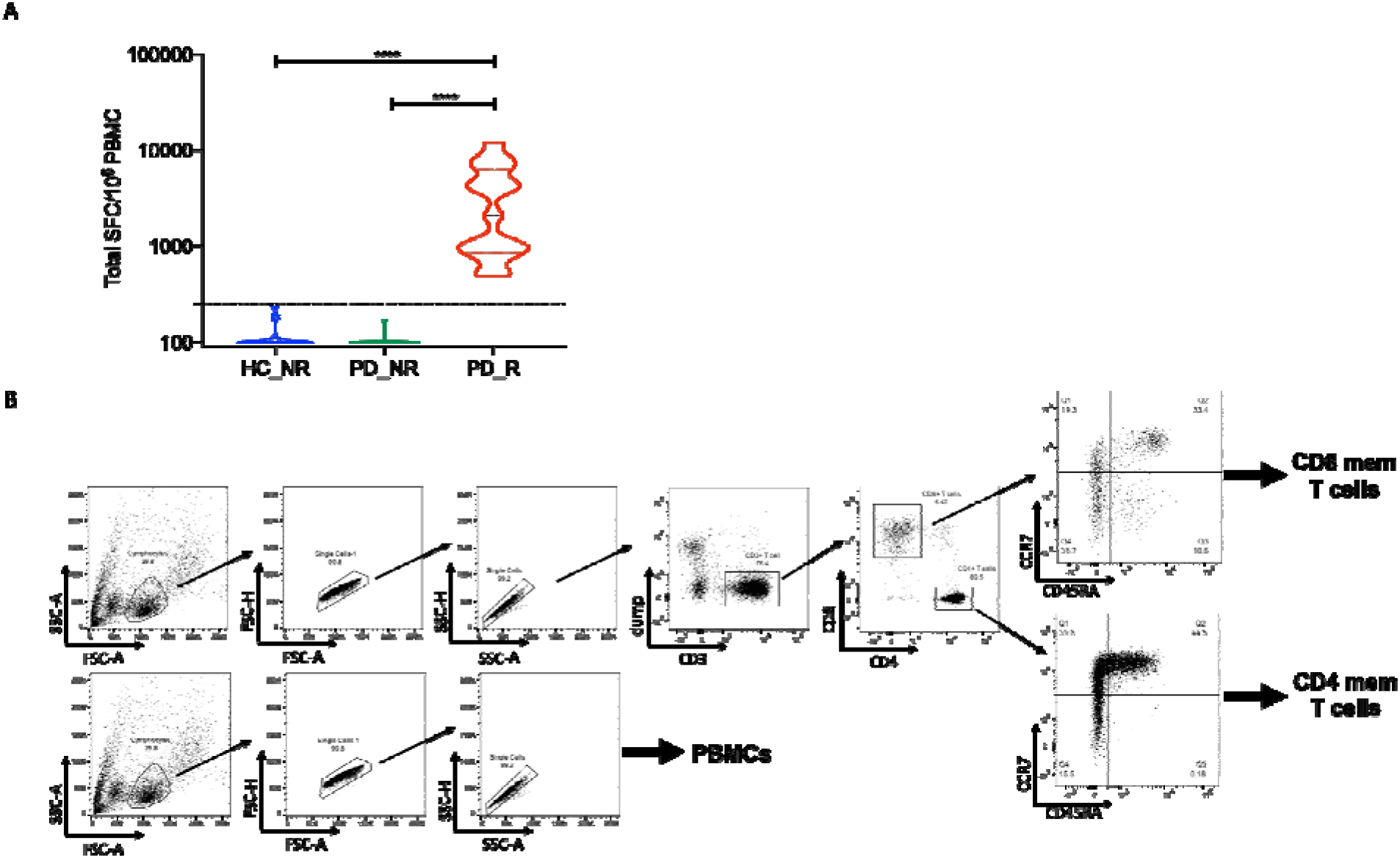
Classification of PD and age-matched HC based on the α-syn T cell response. (A) Violin plot shows the magnitude of T cell response (sum of IFN-γ, IL-5 and IL-10) in HC non-responders (HC_NR) (n=20) PD responders (PD_R) (n=15) and PD non-responders (PD_NR) (n=21). Dotted line denotes the cut off value of 250 SFC. Two -tailed Mann -Whitney, **** p<0.0001 (B) The gating strategy adopted to identify and sort PBMC, CD4 and CD8 memory T cells from PD and HC subjects.

### Transcriptional analysis of PBMC, CD4 and CD8 memory T cells in PD and age-matched HC

We then examined the hypothesis that the circulating peripheral lymphocytes reflect a general inflammatory state associated with early PD. We analyzed PBMC, CD4 and CD8 memory T cells from PD_R, PD_NR, and HC_NR subjects to for specific transcriptomic signatures that might be associated with PD. The low frequency of α-syn-specific CD4 T cells detected in PBMCs in early PD (Lindestam Arlehamn et al., 2020; Sulzer et al., 2017) requires 2-week *in vitro* culture to produce sufficient cells for characterization. CD4 and CD8 memory T cell subsets were identified using CCR7 and CD45RA immunolabel and were sorted based on the gating strategy in **Figure 1B**. Whole PBMC and sorted CD4 and CD8 memory T cell populations were sequenced with the Smart seq protocol (Picelli et al., 2014). To assess whether differences in gene expression could distinguish the groups, we applied Principal Component Analysis (PCA). As expected, the global gene expression profile analyzed by PCA revealed three distinct clusters corresponding to the PBMC, memory CD4 and memory CD8 T cell subsets However, the same analysis did not discriminate between the PBMC, CD4 or CD8 memory T cells from PD and HC subjects (**Supplementary Figure 2A**).

We next performed differential gene expression analysis (DEseq) comparing PD vs. HC_NR to explore PD-specific gene expression signatures of PBMC, CD4 and CD8 memory T cells. Only 26 genes were differentially expressed in PBMC between PD and HC_NR [fold change ≥1.5 (absolute log2 ≥ 0.58) and adjusted p-value <0.05]. Of the 26 genes, only 18 were protein coding; 7 were up-regulated and 11 down-regulated. (**Table 1)**. A total of 11 genes (1 up-regulated and 10 down-regulated; **Table 1**) and 9 genes (4 up-regulated and 5 genes down-regulated; **Table 1**) were differentially expressed protein coding genes in CD4 and CD8 memory T cells, respectively. In conclusion, few genes were differentially expressed at the global level, and we did not identify any specifically molecular pathway that was differentially regulated in PBMC, CD4 or CD8 memory T cells. Moreover, no overlap was observed between the few protein coding genes that were differentially expressed in PD vs. HC_NR, in PBMC, CD4, or CD8 cell subsets (**Supplementary Figure 2B**).

**Table 1.**
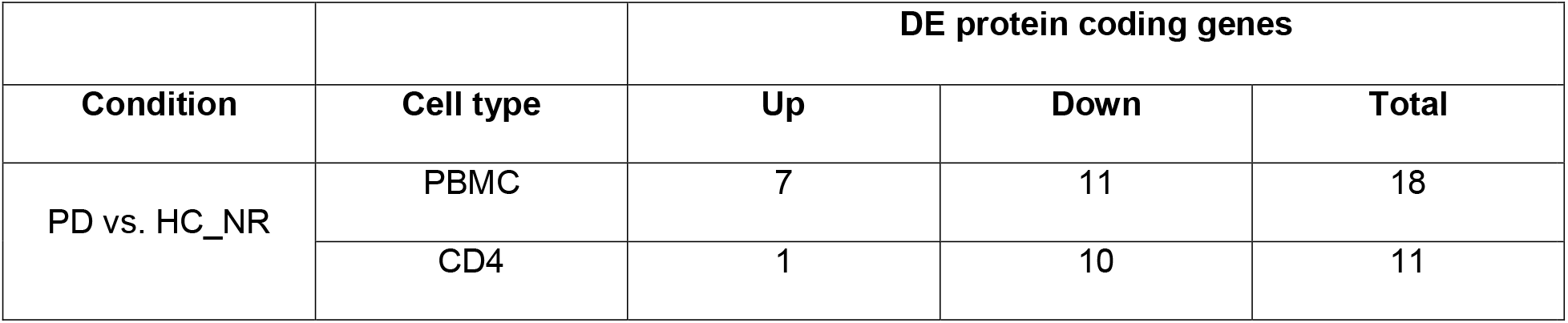
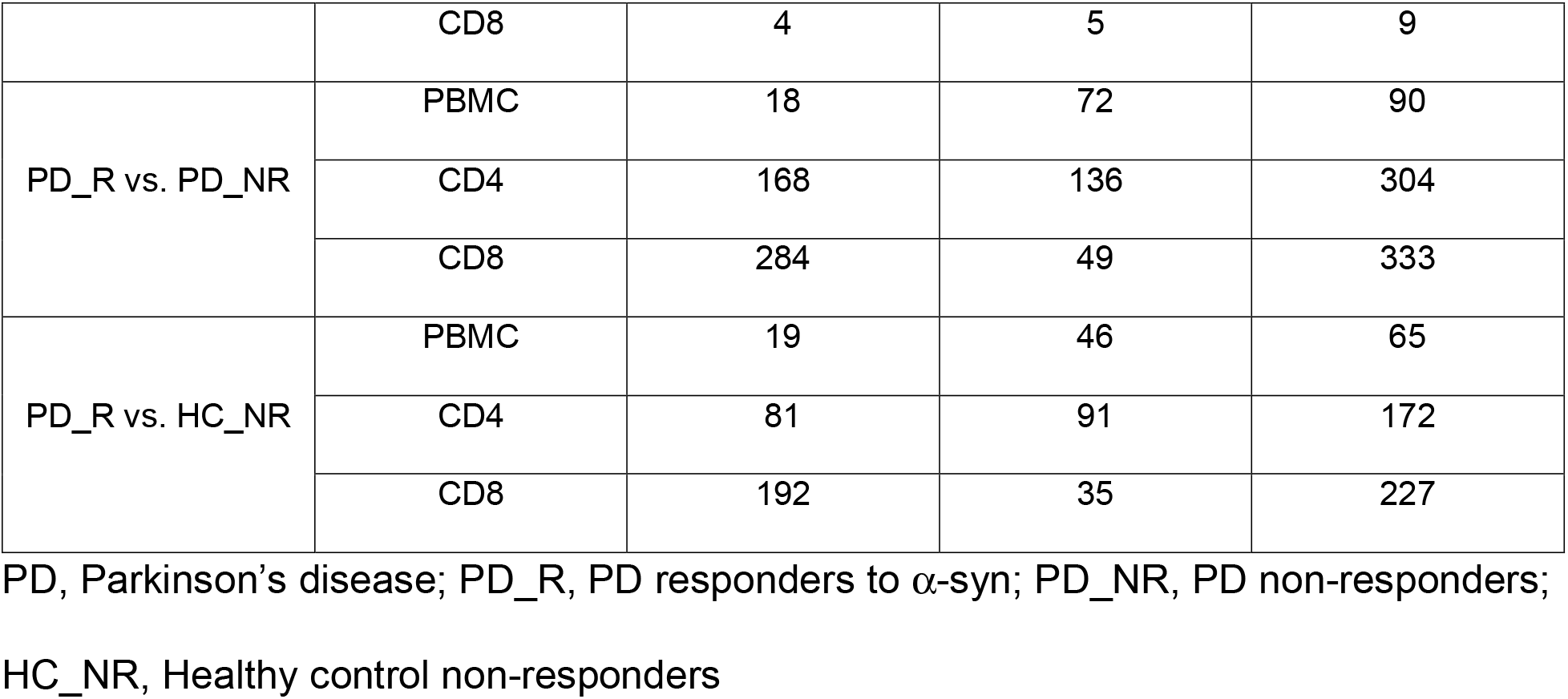
Number of differentially expressed genes in different comparisons.

### Classification of PD subjects based on α-syn-specific T cell reactivity reveals specific gene signatures

Next, we compared the gene expression profiles of PD_R to HC_NR and to PD_NR subjects. We observed a large increase in the number of differentially expressed genes in comparisons of each cell type (PBMC, CD4 and CD8 memory T cells; **Table 1)**. The total number of differentially expressed genes for PBMC between PD_R versus PD_NR and PD_R versus HC_NR was 90 and 65, respectively (**Figure 2A**). Scrutiny of these genes did not reveal any functional enrichment for specific patterns or pathways (**Supplementary Table 1**).

**Figure 2.**
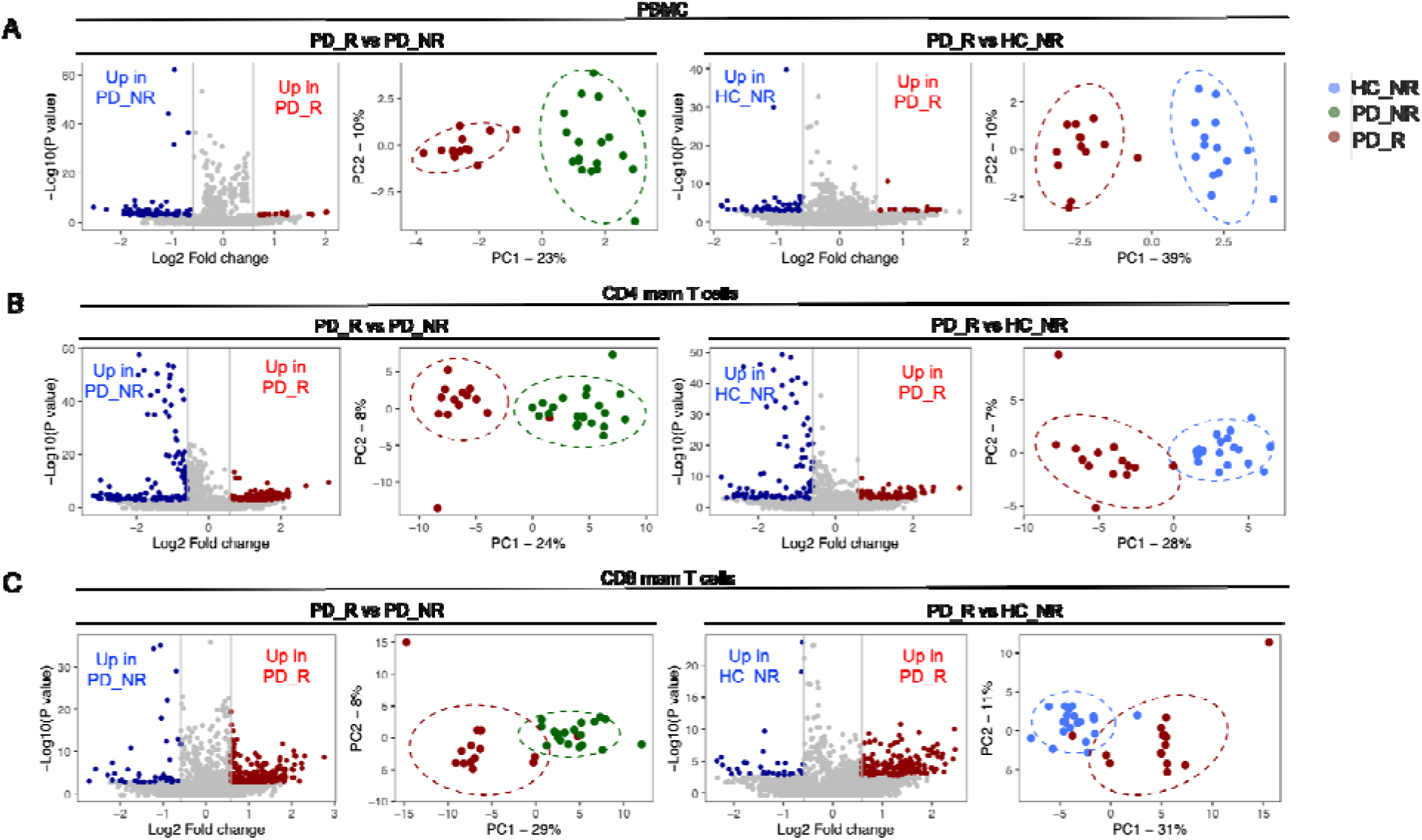
α-syn specific T cell reactivity is associated with a unique gene expression profile. Volcano plots show log_2_ fold change versus -log_10_(P value) for the PD_R (n=15) versus PD_NR (n=21) and PD_R versus HC_NR (n=20) respectively. The subset of genes with an absolute log2 fold change >1.5 and adjusted p-value less than 0.05 were considered significant and are indicated by dotted lines. Red dots of volcano plots indicate protein coding genes upregulated in PD_R and blue dots indicate protein coding genes down-regulated in PD_NR or HC_NR. PCA plots show distinct clusters of PD_R, PD_NR and HC_NR (A) PBMC (B) CD4 memory T cells (C) CD8 memory T cells based on differentially expressed protein coding genes.

In contrast, CD4 and CD8 memory T cells exhibited an intriguing gene signature with an approximately ~2.5 - 4-fold increase in the number of differentially expressed genes between the PD_R and PD_NR groups and between PD_R and HC_NR. PD_R to PD_NR comparison revealed 304 DE genes for CD4 (136 down-regulated and 168 up-regulated; **Figure 2B**), and 333 DE genes for CD8 (49 down-regulated and 284 up-regulated, **Figure 2C, Table 1**). Similarly, comparing PD_R to HC_NR, revealed 172 DE genes for CD4 (91 down-regulated and 81 up-regulated, **Figure 2B**), and 227 DE genes for CD8 (35 down-regulated and 192 up-regulated; **Figure 2C and Table 1)**. As expected, based on the DE genes, the disease groups PD_R, HC_NR, and PD_NR formed distinct clusters (**Figure 2).**There was substantial overlap of DE genes between PD_R vs PD_NR and PD_R vs HC_NR within each cell type, but minimal to no overlap of DE genes across different cell types (**Supplementary Table 1**).

Interestingly, PRKN and LRRK2 genes that are well established to be related to familial PD (Dachsel and Farrer, 2010; Dawson and Dawson, 2010; Di Maio et al., 2018; Li et al., 2014; Mata et al., 2006; Rui et al., 2018; von Coelln et al., 2004), were differentially expressed in CD4 and CD8 memory T cells with both genes down-regulated in CD4 and up-regulated in CD8 memory T cells in PD_R compared to PD_NR and HC_NR respectively (PRKN is up in PD_R vs. PD_NR: LRRK2 is up in PD_R vs HC_NR) indicating that the two cell types play distinct roles in PD-associated T cell autoimmunity. In addition to PRKN, we identified differentially expressed genes including as TFEB and UBAP1L that have been implicated in autophagy and ubiquitination (Settembre et al., 2011) in CD4 memory T cells. Notably, TFEB has been proposed to be a therapeutic target in PD (Decressac and Bjorklund, 2013; Torra et al., 2018; Zhuang et al., 2020).

### Enrichment of PD gene signature in CD4 and CD8 memory T cells

To further characterize the genes differentially expressed in PD_R, HC_NR, and PD_NR, we performed gene set enrichment analysis (GSEA) (Subramanian et al., 2005). To check the enrichment of PD associated gene signature in the differentially expressed genes between PD_R vs. HC_NR and PD_R vs. PD_NR, the DE genes were ranked and compared to an existing gene set “KEGG PARKINSONS DISEASE” that was downloaded from MSigDB in GMT format (Liberzon et al., 2011). As shown in **Figure 3**, a significant enrichment of PD associated genes in PD_R was observed in CD4 and CD8 memory T cells. However, no such enrichment was observed in PBMCs (**Figure 3A**).

**Figure 3.**
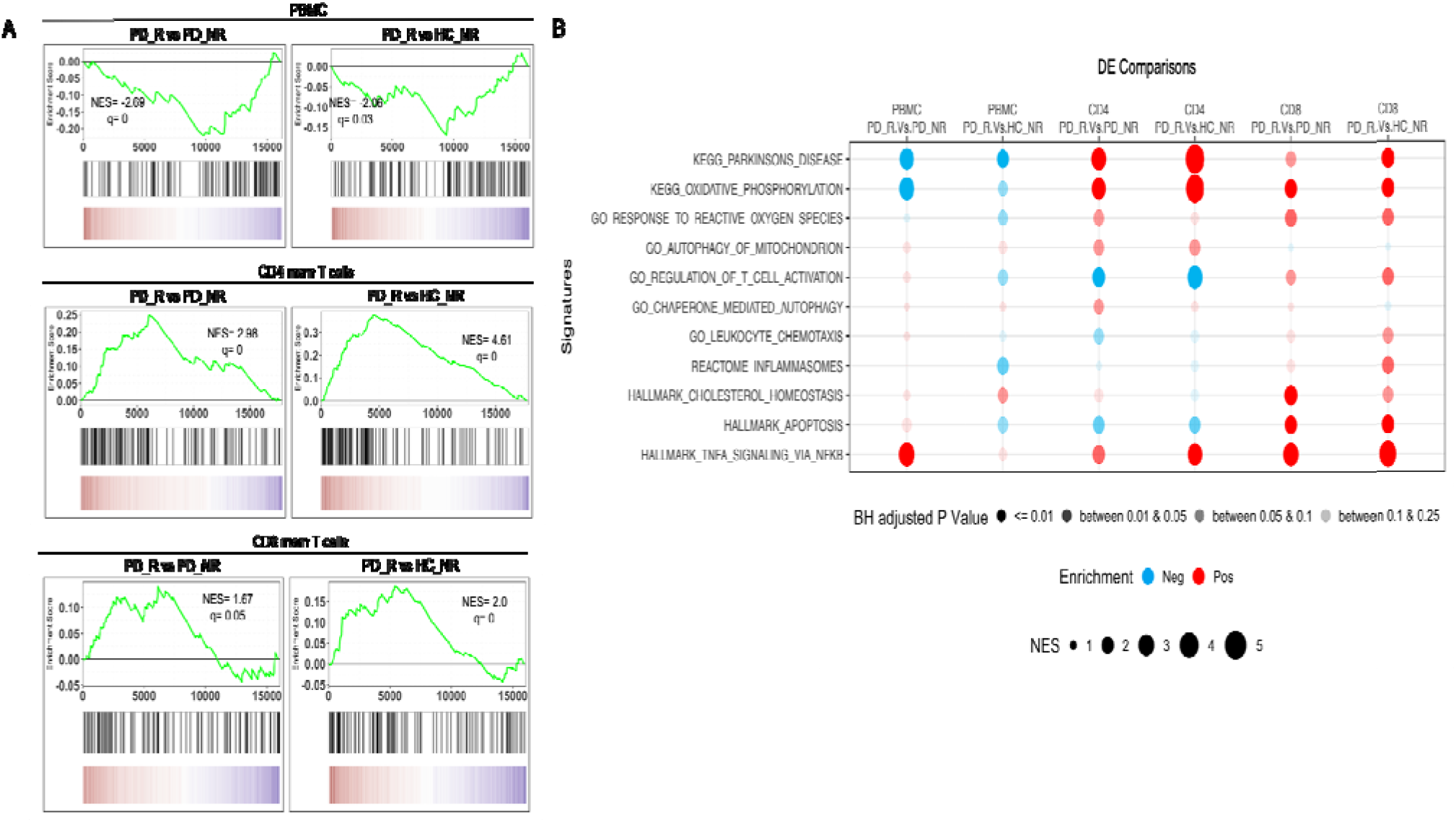
GSEA of the protein coding transcriptome of PD_R vs PD_NR and PD_R vs. HC_NR reveals enrichment of PD associated gene signature in CD4 and CD8 memory T cells. (A) GSEA for the KEGG PD gene set. The y-axis of the plot shows the enrichment score (ES) for the gene set as the analysis moves down the ranked list of genes. The direction of the peak shows the degree to which the gene set is represented at the top or bottom of the ranked list of genes. The black bars on the x-axis show where the genes in the ranked list appear. The red portion at the bottom shows genes upregulated in PD_R and blue portions represents the genes downregulated in PD_R (upregulated in HC_NR or PD_NR). q, false discovery rate; NES, normalized enrichment score. (B) Bubble plot demonstrating the enrichment status of several pathways previously reported to be implicated in PD. The red bubble indicates positive enrichment and blue bubble indicates negative enrichment. The size of the bubble is directly proportional to the normalized enrichment score and the color shade of the bubble is proportional to the adjusted p value, where a darker bubble indicates higher significance than the lighter shade.

We next examined the enrichment of several pathways implicated in PD, including oxidative phosphorylation (Shoffner et al., 1991), oxidative stress (Blesa et al., 2015; Dias et al., 2013; Hemmati-Dinarvand et al., 2019; Hwang, 2013; Jenner, 2003), macroautophagy and chaperone-mediated autophagy (Hou et al., 2020; Lynch-Day et al., 2012; Moors et al., 2017; Wang et al., 2016; Zhang et al., 2012), cholesterol signaling (Jin et al., 2019; Vance, 2012), inflammation (Stojkovska et al., 2015), and TNF signaling (Leal et al., 2013). Interestingly, chemotaxis, apoptosis, cholesterol biosynthesis and inflammation were significantly enriched in CD8 memory T cells and oxidative stress, autophagy of mitochondria and chaperone mediated autophagy were enriched in CD4 memory T cells. Other pathways, such as oxidative phosphorylation and TNF signaling were enriched in both memory T cell subsets (**Figure 3B**).

The results suggest that classifying the PD subjects based on their α-syn T cell reactivity and separately examining memory CD4 and CD8 T cell subsets can detect PD associated gene signatures and identify PD relevant pathways (**Figure 3A-B).** It further suggests that peripheral memory T cell subsets might offer an opportunity to dissect the molecular mechanisms associated with PD pathogenesis. and is consistent with the notion that memory T cells may play a significant role in PD pathogenesis.

### Identification of cell surface and secreted protein targets

Because cell surface expressed or secreted targets are amenable to modulation by monoclonal antibody therapy, we were interested in identifying which of the differentially expressed genes encode surface expressed or secreted products that could be targeted in PD. We performed surfaceome and secretome analysis on the differentially expressed genes between PD_R vs HC_NR and PD_R vs PD_NR in all cell types. For surfaceome analysis, three databases of surface expressing targets (Ashburner et al., 2000; Bausch-Fluck et al., 2018; Bausch-Fluck et al., 2015) were combined and a reference master list of targets that appeared in two out of three databases was generated that comprised of total 1168 targets. For secretome analysis, a reported human secretome database that comprised of 8575 targets was referred (Vathipadiekal et al., 2015). Combining surfaceome and secretome, we identified 133 and 76 targets that were either secretory and/or surface expressed in PD_R vs PD_NR, and PD_R vs HC_NR, respectively, in the CD4 memory T cell subset. We identified 140 and 100 targets in PD_R vs PD_NR, and PD_R vs HC_NR, respectively, in the CD8 memory T cell subset (**Supplementary Table S2**).

### Validation of potential genes of interest

We then selected specific DE genes for validation by flow cytometry based on the availability of commercially available antibodies. Specifically, we validated one DE gene in each cell subset (CCR5 in PBMC; CX3CR1 in memory CD4 subset and CCR1 in memory CD8 subset) at the protein level. The normalized expression count of the genes that were validated is represented in **Supplementary figure 3A**. The protein expression profile of the selected genes largely matched to the gene expression pattern observed by RNAseq analysis (**Supplementary figure 3B**). For example, PBMCs of HC_NR displayed significantly higher expression of CCR5 than PD_R, the CD4 memory subset of PD_NR had higher expression of CX3CR1 than PD_R, and the CD8 memory subset of PD_R had significantly higher expression of CCR1 than PD_NR and HC_NR. Similar trends were observed at the transcriptional and protein levels.

## Discussion

In this study, we show that memory T cells of PD subjects with detectable α-syn responses possess specific mRNA signatures. These signatures are associated with both known genes previously associated with neurological diseases and novel genes. The specific genes and pathways identified that show a significant enrichment of transcriptomic signatures previously associated with PD include oxidative stress, oxidative phosphorylation, autophagy of mitochondria, chaperone-mediated autophagy, cholesterol metabolism, and inflammation. These molecular pathways and the associated genes are known to be dysregulated in PD and are widely thought to accelerate the progression of disease. For instance, dysfunctional autophagic machinery leads to the accumulation of α-syn (Martinez-Vicente et al., 2008) and defective mitochondria (Lee et al., 2012) which in turn can lead to formation of α-syn aggregates or impair energy metabolism and cause oxidative stress. Moreover, the accumulated and misfolded α-syn, a protein normally involved in the regulation of synaptic vesicle exocytosis (Somayaji et al., 2020), causes degeneration of SNpc DA neurons, impairs synapse function (Chung et al., 2009; Ihara et al., 2007; Kahle et al., 2000; Sulzer and Edwards, 2019; Yavich et al., 2006) and affects respiration, morphology, and turnover of mitochondria (Chinta et al., 2010; Choubey et al., 2011; Cole et al., 2008; Devi et al., 2008; Li et al., 2007; Martin et al., 2006; Parihar et al., 2008, 2009), which may be related to display of mitochondrial-derived antigens in PD (Matheoud et al., 2019; McLelland et al., 2014). Additionally, cholesterol metabolism has also been linked to PD (Huang et al., 2019) via a potential role in synaptogenesis. The interplay of implicated pathways suggests that a cascade of several molecular events takes place, resulting in progressive neurodegeneration.

We observed enrichment of reactive inflammasomes in CD8 memory T cell subset of PD responders, but not in their CD4 memory T cell subset, suggesting that PD associated inflammatory signature is cell type specific. We focused on the signatures associated with CD4 and CD8 memory T cells. The focus on T cells is prompted and supported by several reports that imply a T cell-associated inflammatory process (Lindestam Arlehamn et al., 2020; Seo et al., 2020) within the PD prodromal phase and disease progression as well as in animal models (Matheoud et al., 2019). Specific transcriptomic signatures associated with CD4 and CD8 memory T cell compartments have been described in several other pathologies (Burel et al., 2018; Grifoni et al., 2018; Hyrcza et al., 2007; Patil et al., 2018; Tian et al., 2019a; Tian et al., 2019b), including autoimmunity (Hong et al., 2020; Lyons et al., 2010; McKinney et al., 2010), but to our knowledge, this is the first report of such signatures associated with memory T cells in neurodegenerative disease. A key element in our study was to focus on the transcriptional profile of specific purified memory CD4 and CD8 T cell subsets. Should this important aspect not have been considered, most of the differentially expressed genes and associated signatures would have been missed, as exemplified by the fact that very few differentially expressed genes were detected when whole PBMCs were considered.

As recently shown for monocytes, there can be a striking effect of sex on gene expression (Carlisle et al., 2021). The DE genes detected in this study did not suggest sex-specific differences and there was an equal distribution of males and females in the PD-R and PD-NR cohort. Future studies with larger cohorts can provide further insights into the potential differential effects of sex on these signatures.

Transcriptional signatures associated with PD have been reported by several groups based on analysis of samples of neural origin that includes astrocytes, neurons, and brain tissue including substantia nigra (Booth et al., 2019; Keo et al., 2020; Lang et al., 2019; Nido et al., 2020; Sandor et al., 2017). Here, we studied the signatures of T cells isolated from peripheral blood, rather than the CNS, because of the difficulty of accessing the CNS, and importantly, because of the lack of availability of sufficient numbers of T cells available to study in CNS fluids from PD donors and in particular from healthy control subjects (Ransohoff et al., 2003). While future studies might further investigate T cells isolated directly from the CNS, it is known that infiltrating T cells recirculate between the blood and the CNS (Ransohoff et al., 2003; Shechter et al., 2013). To that end, we detected multiple differences in chemokine receptor expression between our PD_R group compared to PD_NR and/or HC_NR. This included reduced CCR5 in PD_R PBMC, as well as a reduction in CX3CR1 signal in PD_R memory CD4 T cells. Interestingly, CCR5 inhibitors have recently been shown to be therapeutic in a non-human primate model of PD (Mondal et al., 2019). As for CX3CR1, its potential role in PD is mainly thought to be mediated through microglia (Angelopoulou et al., 2020); however, the receptor has been shown to define T cell memory populations (Gerlach et al., 2016) which have implications in disease (Yamauchi et al., 2020).

Some of the DE genes found in PBMCs and T cells are implicated in PD pathogenesis. This includes leucine-rich repeat kinase 2 (LRRK2), which is one of the two most common genes associated with familial PD, and is also associated with sporadic PD. It has been noted that LRRK2 is far more highly expressed in immune cells than neurons, and is also linked to Crohn’s disease, an inflammatory bowel disorder, a class of disorders associated with PD (Herrick and Tansey, 2021). LRRK2 expression in PBMCs may be related to regulation of peripheral Type 2 interferon response that lead to dopamine neurodegeneration (Kozina et al., 2018), and its overall expression in T cells and other immune cells can be increased by interferon. In our results, LRRK2 transcript is decreased in PD to levels that are 33% the amount in HC.

Additional genes associated with mechanisms implicated in PD pathogenesis are also differentially expressed in T cells from PD_R subjects, including septin 5 (Son et al., 2005), the GDNF receptor (Sandmark et al., 2018), monoamine oxidase S, aquaporin (Tamtaji et al., 2019), LAMP3 (Liu et al., 2011) which has also been associated with REM sleep disorder (a risk factor for PD (Mufti et al., 2021)), polo-like kinase 1 (Mbefo et al., 2010), and myeloperoxidase (Maki et al., 2019). Most of these genes have been found previously to be expressed in neurons, but here we show for the first time DE of these genes in peripheral cells. Moreover, these and additional DE genes point to the possibility that initiating steps in some PD pathogenic pathways might occur in peripheral immune cells and contribute to multiple hits that lead to the loss of targeted neurons (Raj et al., 2014).

Another key element in our study was a focus on the transcriptional profile of PD subjects that were classified based on their T cell responsiveness to α-syn, which were taken as a proxy for subjects undergoing an ongoing inflammatory autoimmune process. This was a determinant aspect, and if this important aspect not have been considered, most of the differentially expressed genes and associated signatures would have been missed. The classification of subjects based on T cell reactivity of α-syn might be further refined by considering additional antigens other than α-syn that might be also involved in PD pathogenesis (Latorre et al., 2018; Lindestam Arlehamn et al., 2019; Lodygin et al., 2019).

Based on a recently published conceptual model to describe PD pathogenesis (Johnson et al., 2019), factors that contribute to neurodegeneration can be divided into three categories: triggers, facilitators and aggravators. Our study design focused on diagnosed PD patients with established disease, and is therefore likely addressing factors that contribute in disease spread (facilitators) and promote the neurodegenerative process (aggravators). Future studies in at risk categories for PD such as REM sleep disorder cohorts might shed light on RNA signatures associated with disease triggers.

Our data identifies specific genes that could be addressed by therapeutic and diagnostic interventions, including TFEB, SNCA, PARK2 and LRRK2. In a diagnostic setting, detection of alterations in the expression of these genes could contribute to a molecularly-based diagnostic, while in the therapeutic setting, it is possible that early targeting of the same genes by inhibiting or activating their function could delay or terminate disease progression or prevent disease development during the prodromal phase. Supportive of this notion is consistent the observation that anti-TNF treatment (Peter et al., 2018) is associated with lower PD disease incidence.

## Materials and Methods

### Ethics statement

All participants provided written informed consent for participation in the study. Ethical approval was obtained from the Institutional review boards at La Jolla Institute for Immunology (LJI; Protocol Nos: VD-124 and VD-118), Columbia University Medical Center (CUMC; protocol number IRB-AAAQ9714 and AAAS1669), University of California San Diego (UCSD; protocol number 161224), Rush University Medical Center (RUMC; Office of Research Affairs No.16042107-IRB01) and the University of Alabama (UAB; protocol number IRB-300001297).

### Study subjects

For RNAseq, we recruited a total of 56 individuals diagnosed with PD (n=36) and age-matched healthy subjects (n=20) in this study. The subjects were recruited from multiple sites: 32 subjects from Columbia University Medical Center (CUMC) (PD n=26 and HC n=6), 10 subjects from La Jolla Institute for Immunology (LJI) (PD n=4 and HC n=6), 8 subjects from University of California San Diego (UCSD) (PD n=4 and HC n=4), 3 subjects from Rush University Medical Center (RUMC) (PD n=1 and HC n=2), 3 subjects from University of Alabama (UAB) (PD n=1 and HC n=2). For validation cohort, we analyzed 30 subjects: 20 PD and 10 HC. The subjects were recruited from multiple sites: 10 subjects from Columbia University Medical Center (CUMC) (PD n=10), 12 subjects from La Jolla Institute for Immunology (LJI) (PD n=2 and HC n=10), 8 subjects from University of Alabama (UAB) (PD n=8). The characteristics of the enrolled subjects are detailed in **Table 2**.

**Table 2.** Characteristics of the subjects enrolled in the study.

|  | RNAseq Cohort |  |  | Validation cohort |  |  |
| --- | --- | --- | --- | --- | --- | --- |
|  | PD_R | PD_NR | HC_NR | PD_R | PD_NR | HC_NR |
| Total subjects enrolled | 15 | 21 | 20 | 10 | 10 | 10 |
| Median age (range), yr | 70, (49-81) | 66, (44-81) | 67, (50-79) | 67 (44-76) | 65 (46-81) | 52 (22-69) |
| Male, %(n) | 73.3% (11) | 85.7% (18) | 20% (4) | 70% (7) | 70% (7) | 50% (5) |
| Caucasian, % (n) | 88.8 % (32) | 80 % (16) | 20% (4) | 90% (9) | 100% (10) | 50% (5) |
| Median years since diagnosis, (range), yr | 3 (0-12) | 6 (0-16) | NA | 7 (0-12) | 9 (0-20) | NA |
| Median MoCA<sup>a</sup>(range) | 27 (9-30) | 26 (23-30) | NA | 28 (22-30) | 28 (14-30) | NA |
| Median UPDRS<sup>b</sup> (range) | 17 (13-37) | 17 (5-30) | NA | 18 (14-24) | 18 (11-52) | NA |

<sup>a</sup> MoCA collected for n=32 PD patients in the RNAseq cohort and n=17 in the validation cohort
<sup>b</sup> UPDRS collected for n=31 PD patients in the RNAseq cohort and =17 in the validation cohort.

The cohorts were recruited by the clinical core at LJI, by the Parkinson and Other Movement Disorder Center at UCSD, the clinical practice of the UAB Movement Disorders Clinic, and the Movement Disorders Clinic at the department of Neurology at CUMC. PD patients were enrolled on the basis of the following criteria: moderate to advanced PD; 2 of: rest tremor, rigidity, and/or bradykinesia, PD diagnosis at age 45-80, dopaminergic medication benefit, and ability to provide informed consent. The exclusion criteria were atypical parkinsonism or other neurological disorders, history of cancer within past 3 years, autoimmune disease, and chronic immune modulatory therapy. Age matched HC were selected on the basis of age 45-85 and ability to provide written consent. Exclusion criteria were the same as for PD donors and in addition, we excluded self-reported genetic factors. The HC were not screened for prodromal symptoms. The PD patients recruited at RUMC, UAB, CUMC, and UCSD (i.e. not at LJI) all fulfilled the UK Parkinson’s Disease Society Brain Bank criteria for PD.

### Peptides

Peptides were commercially synthesized on a small scale (1 mg/ml) by A&A, LLC (San Diego). A total of 11 peptides of α-syn (Sulzer et al., 2017) were synthesized and then reconstituted in DMSO at a concentration of 40 mg/ml. The individual peptides were then pooled, lyophilized and reconstituted at a concentration of 3.6 mg/ml. The peptide pools were tested at a final concentration of 5 ug/ml.

### PBMC isolation

Venous blood was collected in heparin or EDTA containing blood bags and PBMCs were isolated by density gradient centrifugation using Ficoll-Paque plus (GE #17144003). Whole blood was first spun at 1850 rpm for 15 mins with brakes off to remove plasma. The plasma depleted blood was then diluted with RPMI and 35 ml of blood was gently layered on tubes containing 15 ml Ficoll-Paque plus. The tubes were then centrifuged at 1850 rpm for 25 mins with brakes off. The cells at the interface were collected, washed with RPMI, counted and cryopreserved in 90% v/v FBS and 10 % v/v DMSO and stored in liquid nitrogen.

### Cell sorting

The cryopreserved PBMC were thawed and revived in prewarmed RPMI media supplemented with 5% human serum (Gemini Bio-Products, West Sacramento, CA), 1 % Glutamax (Gibco,Waltham, MA), 1% penicillin/streptomycin (Omega Scientific, Tarzana,CA), and 50 U/ml Benzonase (Millipore Sigma, Burlington, MA). The cells were then counted using haemocytometer, washed with PBS and prepared for staining. The cells at a density of 1 million were first incubated at 4°C with 10% FBS for 10 mins for blocking and then stained with a mixture of the following antibodies: APCef780 conjugated anti-CD4 (clone RPA-T4, eBiosciences), AF700 conjugated anti-CD3 (clone UCHT1, BD Pharmigen), BV650 conjugated anti-CD8a (clone RPA-T8, Biolegend), PECy7 conjugated anti-CD19 (clone HIB19, TONBO), APC conjugated anti-CD14 (clone 61D3, TONBO), PerCPCy5.5 conjugated anti-CCR7 (clone G043H7, Biolegend), PE conjugated anti-CD56 (eBiosciences), FITC conjugated anti-CD25 (clone M-A251, BD Pharmigen), eF450 conjugated anti-CD45RA (clone HI100, eBiosciences) and eF506 live dead aqua dye (eBiosciences) for 30 mins at 4°C. Cells were then washed twice and resuspended in 100 ul PBS for flow cytometric analysis and sorting. The cells were sorted using BD FACSAria- (BD Biosciences) into ice cold Trizol LS reagent (Thermo Fisher Scientific).

### Fluorospot assay

PBMCs were thawed and stimulated for two weeks *in vitro* with α-syn pools. PHA was used as control. Cells were fed with 10 U/ml recombinant IL-2 at an interval of 4 days. After two weeks of culture, T cell responses to a-syn were measured by IFNγ, IL-5 and IL-10 Fluorospot assay. Plates (Mabtech, Nacka Strand, Sweden) were coated overnight at 4°C with an antibody mixture of mouse anti-human IFNγ clone (clone 1-D1K), mouse anti-human IL-5 (clone TRFK5), and mouse anti-human IL-10 (clone 9D7). Briefly, 100,000 cells were plated in each well of the pre-coated Immobilon-FL PVDF 96 well plates (Mabtech), stimulated with the respective antigen at the respective concentration of 5 μg/ml and incubated at 37°C in a humidified CO_2_ incubator for 20-24 hrs. Cells stimulated with α-syn were also stimulated with 10 μg/ml PHA that served as a positive control. In order to assess nonspecific cytokine production, cells were also stimulated with DMSO at the corresponding concentration present in the peptide pools. All conditions were tested in triplicates. After incubation, cells were removed, plates were washed six times with 200 μl PBS/0.05% Tween 20 using an automated plate washer. After washing, 100 μl of an antibody mixture containing IFNγ (7-B6-1-FS-BAM), IL-5 (5A10-WASP), and IL-10 (12G8-biotin) prepared in PBS with 0.1% BSA was added to each well and plates were incubated for 2 hrs at room temperature. The plates were again washed six times as described above and incubated with diluted fluorophores (anti-BAM-490, anti-WASP-640, and SA-550) for 1 hr at room temperature. After incubation, the plates were again washed as described above and incubated with a fluorescence enhancer for 15 min. Finally, the plates were blotted dry and spots were counted by computer-assisted image analysis (AID iSpot, AID Diagnostica GMBH, Strassberg, Germany). The responses were considered positive if they met all three criteria (i) the net spot forming cells per 10^6^ PBMC were ≥ 100 (ii) the stimulation index ≥ 2, and (iii) *p*≤ 0.05 by Student’s t test or Poisson distribution test.

### Smart-seq

PBMC, CD4 and CD8 memory T cells of PD and HC subjects were sorted and total RNA from ~50,000 cells was extracted on a Qiacube using a miRNA easy kit (Qiagen) and quantified using bioanalyzer. Total RNA was amplified according to Smart Seq protocol (Picelli et al., 2014). cDNA was purified using AMPure XP beads. cDNA was used to prepare a standard barcoded sequencing library (Illumina). Samples were sequenced using an Illumina HiSeq2500 to obtain 50-bp single end reads. Samples that failed to be sequenced due to limited sample availability or failed the quality control were eliminated from further sequencing and analysis.

### RNA-seq analysis

The reads that passed Illumina filters were further filtered for reads aligning to tRNA, rRNA, adapter sequences, and spike-in controls. These reads were then aligned to GRCh38 reference genome and Gencode v27 annotations using STAR: v2.6.1 (Dobin et al., 2013). DUST scores were calculated with PRINSEQ Lite (v 0.20.3) (Schmieder and Edwards, 2011) and low-complexity reads (DUST > 4) were removed from the BAM files. The alignment results were parsed via the SAMtools (Li et al., 2009) to generate SAM files. Read counts to each genomic feature were obtained with featureCounts(v1.6.5) (Liao et al., 2014) with default options. After removing absent features (zero counts in all samples), the raw counts were then imported to R/Bioconductor package DESeq2 (v 1.24.0) (Love et al., 2014) to identify differentially expressed genes among samples. Known batch conditions cohort and mapping run id were used in the design formula to correct for unwanted variation in the data. P-values for differential expression were calculated using the Wald test for differences between the base means of two conditions. These P-values are then adjusted for multiple test correction using Benjamini Hochberg algorithm (Benjamini and Hochberg, 1995). We considered genes differentially expressed between two groups of samples when the DESeq2 analysis resulted in an adjusted P-value of < 0.05 and the difference in gene expression was 1.5-fold. The sequences used in this article have been submitted to the Gene Expression Omnibus under accession number GSE174473 (http://www.ncbi.nlm.nih.gov/geo/).

### GSEA

Gene set enrichment analysis was done using the “GseaPreranked” method with “classic” scoring scheme and other default settings. The geneset KEGG PARKINSONS DISEASE was downloaded from MSigDB in GMT format (https://www.gseamsigdb.org/gsea/msigdb/cards/KEGG_PARKINSONS_DISEASE).. Rank files for the DE comparisons of interest were generated by assigning a rank of - log10(p Value) to protein coding genes with log2FoldChange greater than zero and log10(p Value) to genes with log_2_ FoldChange less than zero. The GSEA figures were generated using ggplot2 package in R with gene ranks as the x-axis and enrichment score as the y-axis. The heatmap bar was generated using ggplot with genes ordered by their rank on x-axis and 1 as y-axis. Log_2_FoldChange values were used as the aes color option. scale_colour_gradient2 function was used with a midpoint=0 and other default options.

## Author contributions

RD, JRL, JP performed the experiments. RD, JRL and GW analyzed the data. JRL optimized and performed validation experiments. As.S performed bioinformatic analysis. AF, YX, AWA, DGS, JG, IL and RNA recruited participants and performed clinical evaluations. RD, AS and CSLA wrote the manuscript with substantial edits by DS, BP and other authors. RD, CSLA, AS designed and discussed the data. All authors read, edited, and approved the manuscript.

## Acknowledgements

We would like to thank all the participants for donating samples for this study. We are also grateful to the clinical core for processing the blood samples and flow cytometry core for cell sorting at La Jolla Institute for Immunology. The study is funded by the joint efforts of The Michael J. Fox Foundation for Parkinson’s Research (MJFF) and the Aligning Science Across Parkinson’s (ASAP) initiative, the NIH, and the JPB Foundation. MJFF administers the grant ASAP-000375 on behalf of ASAP and itself.

