## Supplemental table 2 for "Transcriptional analysis of peripheral memory T cells reveals Parkinson’s disease-specific gene signatures"

**Supplementary Table 2: Surface expressing and secretory targets in different comparisons:** Genes in bold are both surface and secretory targets, unbolded genes are secretory, italicized genes are surface expressing targets. DE: Differentially expressed genes, SE: surface expressing/secretory target gene.

| | PBMC | | | CD4 memory T cells | | | CD8 memory T cells | | |
|---|---|---|---|---|---|---|---|---|---|
| **Comparison** | **PD vs HC** | **PD_R vs PD_NR** | **PD_R vs HC_NR** | **PD vs HC** | **PD_R vs PD_NR** | **PD_R vs HC_NR** | **PD vs HC** | **PD_R vs PD_NR** | **PD_R vs HC_NR** |
| **DE genes** | 26 | 132 | 101 | 16 | 503 | 260 | 14 | 494 | 356 |
| **Protein coding** | 18 | 90 | 65 | 11 | 304 | 172 | 9 | 333 | 227 |
| **SE genes** | 9 | 39 | 25 | 4 | 133 | 76 | 3 | 140 | 100 |
| **Down regulated Genes** | **DCHS1**<br>TMTC1<br>WDR5B<br>CACNA1F | POPDC2<br>GPR171<br>AATK<br>XPNPEP2<br>P2RY6<br>NAPSA<br>**SLC45A4**<br>PRPF40B<br>FAM173B<br>**SLC22A16**<br>DKK3<br>NUDT6<br>BIK<br>C1QC<br>PLCD3<br>MADCAM1<br>HAS1<br>**HFE**<br>ORMDL2<br>EMID1<br>C17orf80<br>STK32B<br>**SLC22A23**<br>**CCR5**<br>**CXCR1**<br>CLIP3<br>LRRC3<br>F2RL3<br>SLCO5A1<br>HIGD1A<br>*MRC1* | P2RY6<br>EPHX3<br>**NRP2**<br>**ACE**<br>**SLC22A16**<br>LRRC3<br>PRSS27<br>**HFE**<br>FFAR1<br>PRPF40B<br>**CCR5**<br>FUT2<br>SDCBP2<br>TMTC1<br>**DCHS1**<br>CTRC<br>COL4A2 | ZACN<br>GREM2<br>CYP2U1 | CALHM2<br>MAMDC4<br>HS6ST1<br>PEX26<br>CDC42EP<br>2<br>**CD36**<br>**CD8A**<br>MMP25<br>CISH<br>TNFAIP2<br>FGR<br>ACAN<br>ABCC3<br>RAB6B<br>TMEM201<br>IGFBP6<br>**CX3CR1**<br>F2RL2<br>AP2A1<br>FBLN2<br>DHCR24<br>KCNN3<br>**ITGB4**<br>SPINK4<br>**MFSD8**<br>ST6GALN<br>AC6<br>TBC1D8<br>FCGBP<br>**CSF3R** | SYK<br>CDA<br>LAT2<br>KCNH4<br>DHCR24<br>**CCR3**<br>NPHP4<br>F2RL2<br>COL16A1<br>CD300LB<br>**MSLN**<br>AP2A1<br>RAB26<br>IGFBP6<br>RNF152<br>ACAN<br>CPB2<br>**SLC15A2**<br>CYTL1<br>ZACN<br>LRRK2<br>TREML1<br>WIPI1<br>SEMA3G<br>**CD36**<br>SLC7A8<br>VSIG4<br>ABCC3<br>IL22<br>TOM1L1<br>FXYD6 | SEMA6C<br>*PODXL* | KDELR2<br>DMXL2<br>KCNQ4<br>CNIH2<br>**IL10RB**<br>TMEM203<br>BAIAP2L1<br>MMP17<br>TACR2<br>**TMEM179B**<br>VAMP4<br>AMACR<br>RNF5<br>HIST1H4I<br>EPS8L1<br>CDC42EP2<br>HCN2<br>ZNRF3<br>PRKD2<br>PAQR4<br>KDELR1 | PRKD2<br>ACRBP<br>KCNQ4<br>CNIH2<br>CD320<br>GIPR<br>SLC25A42<br>UPF3A<br>MAOA<br>BAIAP2L1<br>SLC25A17<br>PAQR4<br>SYT6<br>C14orf132<br>PELP1<br>VAMP4<br>REG4<br>SEMA6C<br>EPS8L1<br>*PODXL* |

| | | | | | | |
|---|---|---|---|---|---|---|
| | | *F2RL3* | | | XPNPEP2<br>**SIGLEC7**<br>SRC<br>WIPI1<br>LAT2<br>PSKH1<br>RAB3D<br>CDA<br>**SELPLG**<br>COL16A1<br>TOM1L1<br>**HHLA2**<br>KCNH4<br>RAB1B<br>RNF152<br>CPB2<br>CD300LB<br>SGCA<br>FFAR3<br>PBXIP1<br>**LRFN2**<br>SLC7A8<br>TAP1<br>CHRNA10<br>**RET**<br>EXOC3L2<br>IL22<br>ST3GAL6<br>**SLC15A2**<br>GPR153<br>MS4A14<br>KIRREL2<br>LRRK2<br>SLC30A8<br>MPO<br>OLFML2B<br>LYPD4<br>**IGDCC4**<br>GML<br>FGFBP1 | FFAR3<br>GBGT1<br>STRC<br>SLC35F3<br>SCARA3<br>MCOLN3<br>**LAMP3**<br>ZNF532<br>TRPM2<br>MTUS1<br>PDIA5<br>AIG1<br>INVS<br>TMEM97<br>H6PD<br>FAM19A2<br>CYP2S1<br>LEAP2<br>CASK<br>SARNP<br>**PTGIR**<br>**CSF3R**<br>SLC25A3<br>0<br>RAB6B<br>TMEM201<br>C2CD2<br>CHRNA10<br>NAPA |

| | | | | | | | | | |
|---|---|---|---|---|---|---|---|---|---|
| **Upregulated genes** | CLEC4F<br>AP3B2<br>ITPRIPL1<br>EIF1AY<br>CYP2F1 | TPR<br>EMILIN2<br>ZNRF3<br>SPOCD1<br>CTDSPL<br>EYS<br>GCNT7 | PCBP2<br>CERK<br>**LPAR1**<br>EIF1AY<br>PEX3<br>PDPR<br>**CRIM1**<br>CYP2F1 | SOX12 | GYPE<br>RASD1<br>**CD180**<br>B3GNT8<br>STRC<br>SLC7A10<br>TOMM5<br>RNASEL<br>ADCK1<br>FUT11<br>INVS<br>**CSF2RB**<br>EPHX1<br>HIST1H4H<br>SLC25A19<br>CYP2S1<br>TRPM2<br>MAN1C1<br>PEX11G<br>NDUFC1<br>**CELSR2**<br>CREB3L4<br>MEGF6<br>MPP5<br>TMEM97<br>ECE2<br>GCNT2<br>SVIL<br>SARNP<br>PDIA5<br>**ADAM22**<br>ALS2<br>MGP<br>H6PD<br>PIGK<br>DENND1A<br>BCL2<br>CHMP5<br>CRB3<br>KCNQ1<br>**FAM171A1** | AQP9<br>ECE2<br>INSL4<br>TOMM5<br>SERPINH1<br>HIST1H4H<br>GDF11<br>ABCD2<br>CRB3<br>**CELSR2**<br>MPHOSPH9<br>**TNFRSF11A**<br>GALNT1<br>APOL1<br>DCST1<br>BAIAP2<br>SOX12 | **GPM6B** | MSRB3<br>LTC4S<br>METTL7A<br>OVGP1<br>MAPK15<br>**SLC19A1**<br>MTX2<br>COL1A1<br>MBOAT2<br>SORD<br>HSD17B7<br>SLC4A8<br>**CDH5**<br>GNG12<br>**ANO6**<br>MAMLD1<br>C8G<br>ANKRD44<br>PLEKHB1<br>TSSK4<br>**CALCRL**<br>MGLL<br>MFSD6L<br>QPCT<br>NDFIP2<br>MS4A6A<br>MARCO<br>CLCN1<br>SEMA3B<br>C2CD2<br>GPRIN1<br>CLN6<br>PLXDC2<br>KCNH3<br>APOO<br>ZNF599<br>B3GALT2<br>FBXO2<br>GLT1D1<br>LAPTM4B<br>**CTLA4** | TMPRSS2<br>**GFRA2**<br>MFSD6L<br>LRRK2<br>SRC<br>RGMB<br>B4GALNT1<br>LRRC3<br>OVGP1<br>**CCR1**<br>**SLC19A1**<br>S100A8<br>G0S2<br>GNG12<br>SH2B2<br>FPR1<br>**PLXNA4**<br>LHFPL2<br>ATP6V0A1<br>ZDHHC14<br>C2CD2<br>HSD17B6<br>CD300C<br>**MEGF8**<br>LYZ<br>GRASP<br>MAPK15<br>BTK<br>PLOD3<br>**NTSR1**<br>LAPTM4B<br>TIAM1<br>PGLYRP2<br>CHRNA10<br>**CCR8**<br>KCNH3<br>AIF1<br>LGALS3BP<br>PRRG4<br>**SV2A**<br>FAR2 |

| | | | | | | | | | |
|---|---|---|---|---|---|---|---|---|---|
| | | | | | ZNF532<br>MTUS1<br>ISM1<br>DYNC2H1<br>NLRP2<br>AP2A2<br>SERPINH1<br>DCST1<br>APOL1<br>AIG1<br>**LAMP3**<br>ADCK5<br>**IL6R**<br>CASK<br>MPHOSPH9<br>**ALCAM**<br>FAM19A2<br>ABCD2<br>PXMP2<br>CNKSR2<br>GDF11<br>BAIAP2<br>LEAP2 | | | **CCR1**<br>**CXADR**<br>FITM2<br>RASGRF2<br>MAP2K6<br>DRD3<br>S100A8<br>TPCN1<br>BCL7A<br>IL17C<br>ZDHHC14<br>LAT2<br>KIAA0319L<br>**NOTCH4**<br>**NTSR1**<br>**GFRA2**<br>HMOX1<br>RALGPS2<br>RASL11A<br>B3GNT5<br>PLOD3<br>RGMB<br>GCA<br>CHRNA10<br>LGALS3BP<br>GPR75<br>DHRS12<br>CD300C<br>HOMER1<br>HSD17B6<br>SRC<br>FCN1<br>COL9A2<br>LTK<br>ACSM3<br>TNFSF13B<br>SPON1<br>SCRIB<br>IL12A<br>SVIL<br>FXYD2<br>BIK | LTK<br>RALGPS2<br>SVIL<br>**CALCRL**<br>PCYOX1L<br>TSSK4<br>MSRB3<br>CYB561D1<br>SLC24A4<br>**EPHB3**<br>IL10<br>TNFSF13B<br>MGLL<br>WDR44<br>LAT2<br>**GPM6B**<br>HSPA13<br>**SEMA6B**<br>**CNTNAP1**<br>IL17C<br>**CTLA4**<br>PPL<br>HOOK1<br>DDN<br>PHLPP1<br>PGAP3<br>JUP<br>SPRN<br>FCN1<br>**SLC1A2**<br>RASGRF2<br>FITM2<br>TMEM170B<br>S100A9<br>**TTYH3**<br>PRSS22<br>CFP<br>SLC38A7<br>HMOX1 |

| | | | | | | | | |
|---|---|---|---|---|---|---|---|---|
| | | | | | | | | ALOX5<br>TIAM1<br>AIF1<br>**DPP4**<br>CDHR1<br>ENOX2<br>**SLC1A2**<br>ACTN1<br>F5<br>CYB561D1<br>PGAP3<br>IMPACT<br>**PLXNA4**<br>DUSP6<br>HSPA13<br>S100A9<br>CHIC1<br>TMEM170B<br>SARNP<br>SEMA6A<br>KCNQ5<br>LYZ<br>**SV2A**<br>SLC38A7<br>LGMN<br>ATP8B3<br>TBXAS1<br>IFT140<br>CCDC136<br>LYSMD4<br>**TTYH3**<br>SPRN<br>CFP<br>JSRP1<br>**PTPRS**<br>PPL |
