## Supplemental figures for "Transcriptional analysis of peripheral memory T cells reveals Parkinson’s disease-specific gene signatures"

### Supplementary figures

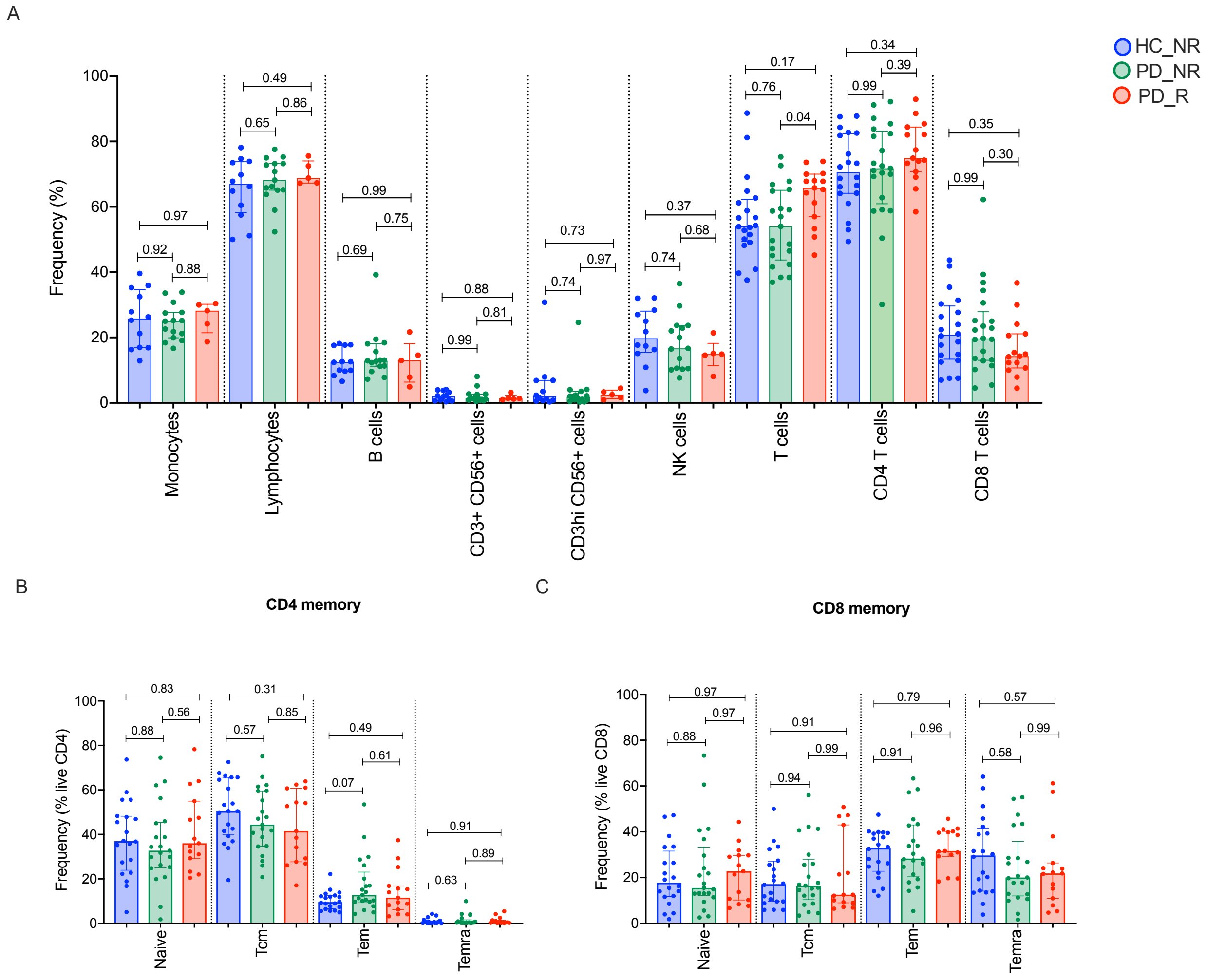


**Figure S1. Relative frequency of different cell subsets in HC_NR, PD_NR and PD_R**. (A). Frequency of major PBMC subsets in HC_NR (blue bar and circles), PD_NR (green bars and circles) and PD_R (red bars and circles) (B) CD4 memory and (C) CD8 memory T cells were further evaluated for frequency of naïve, effector memory ($T_{em}$), central memory ($T_{cm}$) and $T_{EMRA}$ populations. Each point represents a donor. Median ± interquartile range is displayed. Anova with multiple comparison Tukey correction.

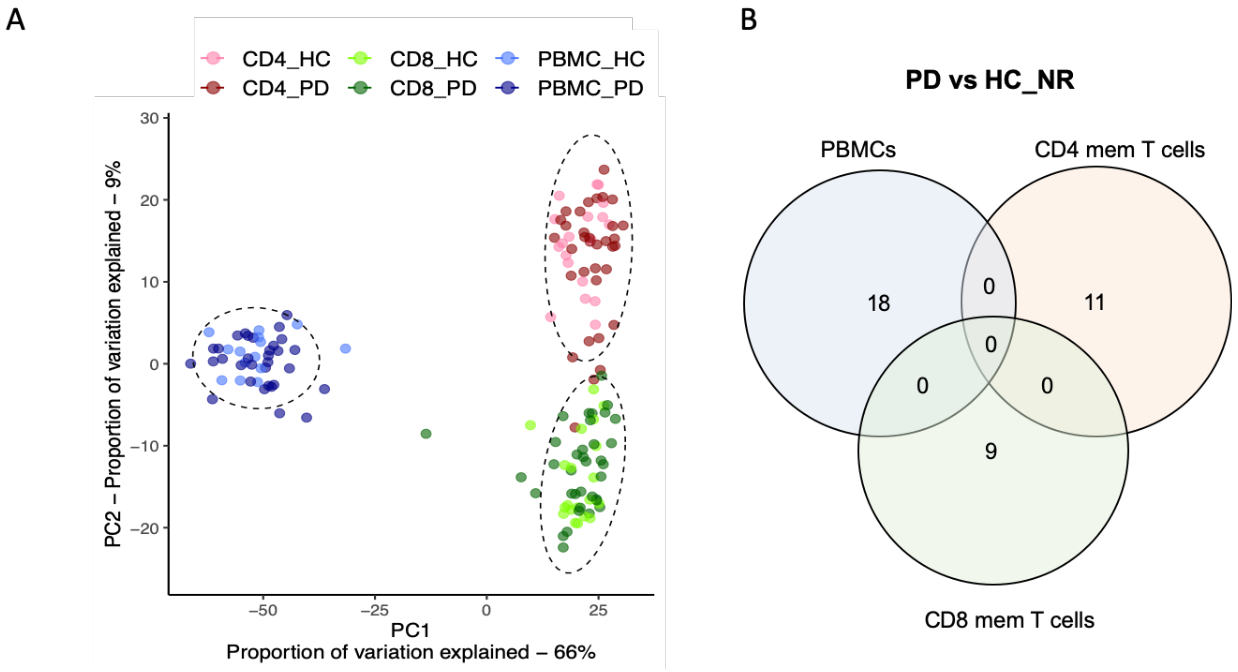


**Figure S2**. **Comparison of PD vs HC in PBMCs, CD4 and CD8 memory T cells** (A) PCA plot demonstrating distinct profile of PBMCs, CD4 and CD8 memory T cells and no separation between PD and HC_NR in either cell type. (B) Venn diagram demonstrating the overlap between PBMC, CD4 and CD8 memory T cells.

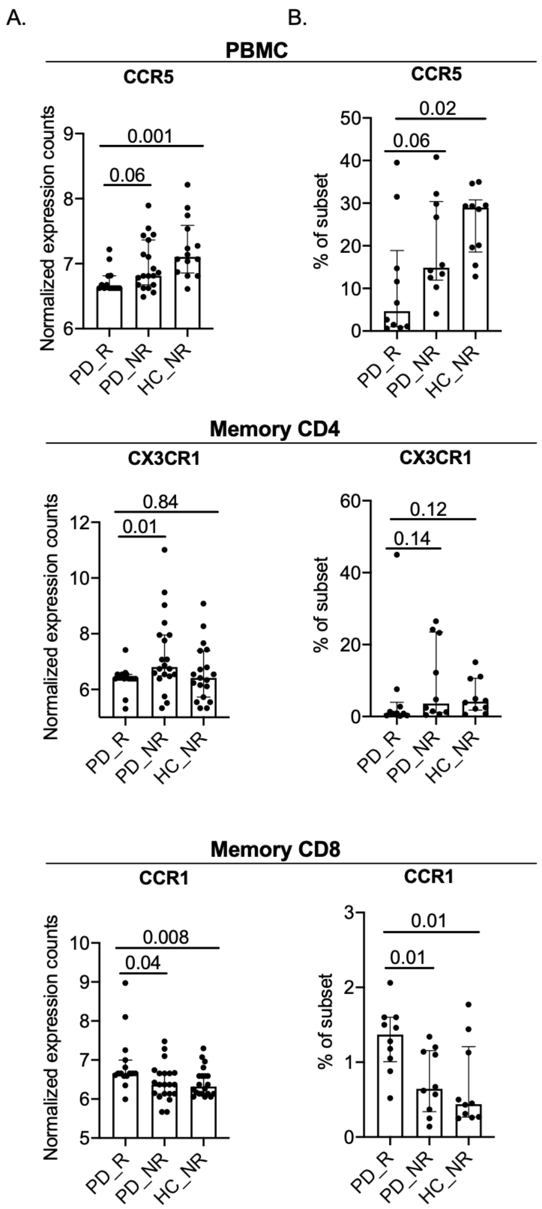


**Figure S3. Gene expression profile of specific DE genes in PBMC, CD4 memory and CD8 memory cell types.** (A) Gene expression values of CCR5, CX3CR1, and CCR1 in counts normalized by sequencing depth calculated by DEseq2 package. (B) Protein expression as percent frequency of subset measured using flow cytometry. Median interquartile range is shown. Two-tailed Mann-Whitney test.
